# Rescue of aberrant huntingtin palmitoylation ameliorates mutant huntingtin-induced toxicity

**DOI:** 10.1101/2021.03.26.437210

**Authors:** Fanny L. Lemarié, Nicholas S. Caron, Shaun S. Sanders, Mandi E. Schmidt, Yen T.N. Nguyen, Seunghyun Ko, Xiaohong Xu, Mahmoud A. Pouladi, Dale D.O. Martin, Michael R. Hayden

## Abstract

Huntington disease (HD) is a neurodegenerative disorder caused by a CAG expansion in the *HTT* gene that codes for an elongated polyglutamine tract in the huntingtin (HTT) protein. HTT is subject to multiple post-translational modifications (PTMs) that regulate its cellular function. Mutating specific PTM sites within mutant HTT (mHTT) in HD mouse models can modulate disease phenotypes, highlighting the key role of HTT PTMs in the pathogenesis of HD. These findings have led to increased interest in developing small molecules to modulate HTT PTMs in order to decrease mHTT toxicity. However, the therapeutic efficacy of pharmacological modulation of HTT PTMs in preclinical HD models remains largely unknown. HTT is palmitoylated at cysteine 214 by the huntingtin-interacting protein 14 (HIP14 or ZDHHC17) and 14-like (HIP14L or ZDHHC13) acyltransferases. Here, we assessed if HTT palmitoylation should be regarded as a therapeutic target to treat HD by (1) investigating palmitoylation dysregulation in rodent and human HD model systems, (2) measuring the impact of mHTT-lowering therapy on brain palmitoylation, and (3) evaluating if HTT palmitoylation can be pharmacologically modulate. We show that palmitoylation of mHTT and some HIP14/HIP14L-substrates is decreased early in multiple HD mouse models, and that aging further reduces HTT palmitoylation. Lowering mHTT in the brain of YAC128 mice is not sufficient to rescue aberrant palmitoylation. However, we demonstrate that mHTT palmitoylation can be normalized in COS-7 cells, in YAC128 cortico-striatal primary neurons and HD patient-derived lymphoblasts using an acyl-protein thioesterase (APT) inhibitor. Moreover, we show that modulating palmitoylation reduces mHTT aggregation and mHTT-induced cytotoxicity in COS-7 cells and YAC128 neurons.

**Highlights:**

- Palmitoylation of mHTT is reduced in multiple transgenic HD mouse models
- HTT palmitoylation decreases with increasing polyQ length in HD patient cells
- mHTT-lowering in mouse brains does not rescue aberrant palmitoylation
- mHTT palmitoylation in HD patient-derived cells can be rescued via APT inhibition
- Promoting palmitoylation reduces mHTT aggregation and cytotoxicity *in vitro*

## 1. Introduction

Huntington disease (HD) is an inherited neurodegenerative disorder characterized by involuntary movements, behavioural changes and cognitive impairment. There are currently no therapies available to influence its inexorable progression (Caron et al., 2018; Evans et al., 2013; Fisher and Hayden, 2014; Myers, 2004). The genetic cause of HD is an expanded CAG trinucleotide repeat (>35) in the *HTT* gene leading to an abnormally long polyglutamine (polyQ) tract in the huntingtin (HTT) protein.

HTT interacts with numerous protein partners and is involved in many cellular processes, including endocytosis, vesicle/organelle transport and recycling, autophagy, and DNA transcription (Maiuri et al., 2017; Ochaba et al., 2014; Saudou and Humbert, 2016). HTT is subject to numerous post-translational modifications (PTMs), including phosphorylation (Aiken et al., 2009; Humbert et al., 2002; Schilling et al., 2006), acetylation (Jeong et al., 2009), ubiquitination (Kalchman et al., 1996), SUMOylation (Steffan et al., 2004), proteolysis (Graham et al., 2006), and fatty acylation (palmitoylation and myristoylation) (Martin et al., 2018, 2014; Yanai et al., 2006). These PTMs regulate subcellular localization, protein-protein interactions, aggregation and clearance which modulates the toxicity of mutant HTT (mHTT) (Atwal et al., 2011, 2007; Cariulo et al., 2017; Caron et al., 2013; Kratter et al., 2016; Lontay et al., 2020).

In the presence of the HD mutation, levels of several key PTMs are significantly altered *in vitro* and *in vivo* (Cariulo et al., 2020, 2017; Warby et al., 2005). Introducing mutations at specific PTM sites of mHTT has been shown to modulate disease phenotypes in various HD models. Blocking proteolysis of mHTT at aspartic acid 586 (D586) prevents striatal neurodegeneration in the YAC128 HD mouse model (Graham et al., 2006).

Blocking phosphorylation at serine residues 13/16 or 421 (S13/16A, S421A) in the BACHD mouse model and HD human induced pluripotent stem cell (hiPSC)-derived neural cells exacerbates disease phenotypes. In contrast, mimicking phosphorylation at these residues (S13/16D, S421D) prevents neurodegeneration in these same models (Gu et al., 2009; Kratter et al., 2016; Xu et al., 2020). These findings highlight the key role of HTT PTMs in the pathogenesis of HD and have stimulated interest in the development of drugs to modulate HTT PTMs in order to decrease mHTT levels and toxicity. To date, most research has focused on normalizing HTT phosphorylation at S13 and S16, and preventing mHTT proteolysis (Aharony et al., 2015; Atwal et al., 2011; Bowie et al., 2018).

Palmitoylation refers to the addition of a palmitic acid (C16:0) onto a cysteine residue via a thioester bond. This process is catalyzed by a family of palmitoyl acyltransferases (PATs). Palmitoylation increases the hydrophobicity of proteins, and therefore plays a key role in protein trafficking, stability, membrane association and protein-protein interactions (Fukata and Fukata, 2010). It is the most common protein-lipid modification in the brain (Smotrys and Linder, 2004). Importantly, it is reversible and allows for dynamic regulation of protein functions at the synapse, thus playing an important role in neurodevelopmental processes and in neuronal survival (Globa and Bamji, 2017; Hubalkova et al., 2021; Matt et al., 2019; Mukai et al., 2008). Disruption of protein palmitoylation has been implicated in the pathogenesis of several neurological disorders, including Alzheimer’s disease, schizophrenia and intellectual disability (Cho and Park, 2016; Sanders et al., 2015b).

We first described that HTT is palmitoylated at cysteine 214 (C214), a reaction catalyzed by the palmitoyl acyltransferases ZDHHC17 and ZDHHC13, also known as huntingtin-interacting protein 14 and 14-like (HIP14 and HIP14L) (Yanai et al., 2006). In the presence of the HD mutation, mHTT was found to be hypo-palmitoylated in the brain of the YAC128 HD mouse model (Yanai et al., 2006). Transient expression of mHTT carrying a palmitoylation-resistant mutation (C214 to serine, C214S) in immortalized cell lines and primary neurons led to increased aggregates and nuclear inclusion formation, increased cell death and increased susceptibility to excitotoxicity (Yanai et al., 2006). Autopalmitoylation of HIP14 and HIP14L, and palmitoylation of many of their synaptic substrates were also downregulated in the YAC128 HD mouse brains or in the brains of mice lacking one *Htt* allele (*Htt*^+/−^) (Huang et al., 2011; Singaraja et al., 2011). Moreover, mouse models deficient in either HIP14 (*Hip14*^-/-^) or HIP14L (*Hip14l*^-/-^) develop neuropathological deficits that are reminiscent of HD (Sutton et al., 2013), and loss of both *Hip14* and *Hip14l* leads to embryonic lethality (Sanders et al., 2016, 2015a). Collectively, these findings link defects in fatty acylation with neuronal dysfunction and neurodegeneration in HD. However, the effect of palmitoylation on HTT function and the role that loss of HTT palmitoylation plays in the pathogenesis of HD are still not fully elucidated.

In this study, we investigate whether palmitoylation is dysregulated in HD mouse models and human HD samples. We also evaluate whether pharmacological modulation of mHTT palmitoylation represents a potential therapeutic target for the treatment of HD.

## 2. Materials and methods

### 2.1. Materials

#### 2.1.1. Reagents and chemicals

**Table 1** lists the reagents and chemicals used for this study along with the application, manufacturer and catalogue number.

**Table 1.**
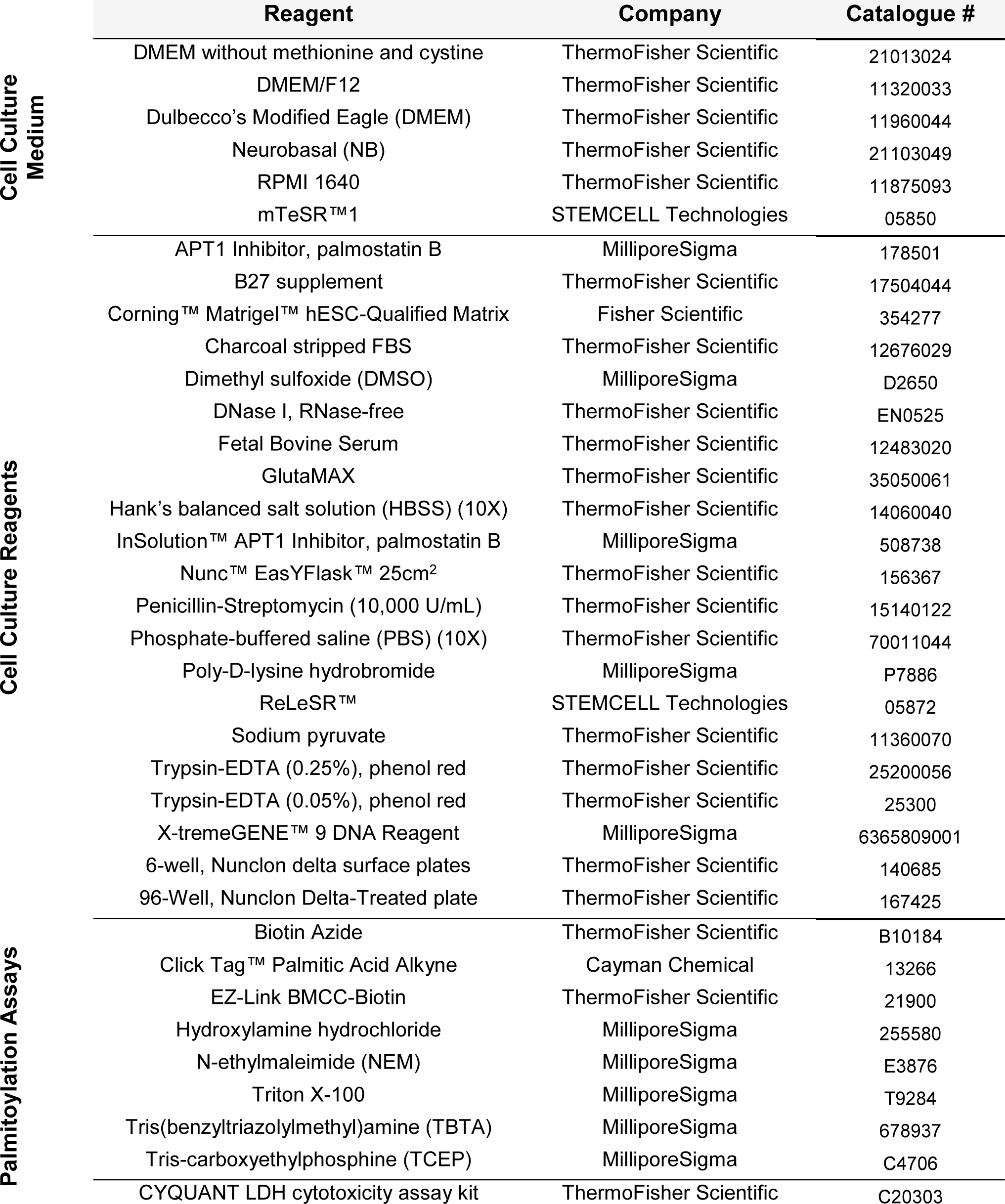
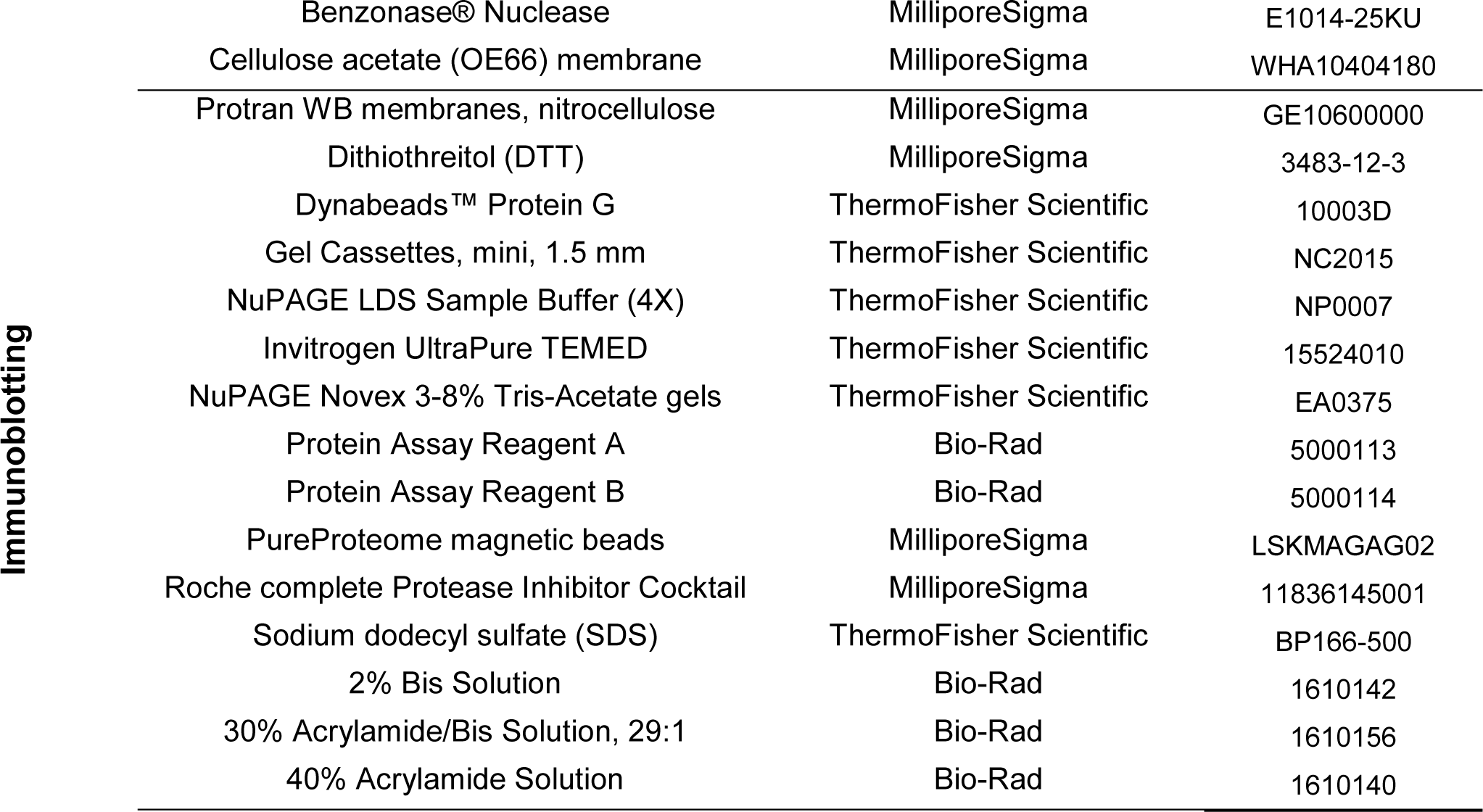
List of reagents and chemicals used and application.

#### 2.1.2. Antibodies

**Table 2** displays the list of antibodies used along with the application, concentrations and experimental conditions.

**Table 2.** List of antibodies used and application.

| Target (clone) | Species | Type | Supplier | Catalogue AB Registry Reference | Application | Dilution | Volume* |
| --- | --- | --- | --- | --- | --- | --- | --- |
| HTT (1HU-4C8) | Mouse | Monoclonal | Sigma-Aldrich | MAB2166 AB_2123255 | WB | 1:1,000 | - |
|  |  |  |  |  | IP | - | 2 µL |
| HTT (D7F7) | Rabbit | Monoclonal | Cell Signaling Technology | 5656 AB_10827977 | WB | 1:1,000 | - |
| HTT (BKP1/HD46) | Rabbit | Polyclonal | in-house | (Kalchman et al., 1997) | IP | - | 2 µL |
|  |  |  |  |  | WB | 1:400 | - |
| HTT | Rabbit | Polyclonal | Sigma-Aldrich | H7540 AB_1840946 | WB | 1:1,000 | - |
|  |  |  |  |  | IP | - | 2 µL |
| Streptavidin | - | Conjugated to AF-680 | Invitrogen | S-32358 | WB | 1:5,000 | - |
| PSD95 (7E3-1B8) | Mouse | Monoclonal | Invitrogen | MA1-046 AB_2092361 | WB | 1:1,000 | - |
| PSD95 | Rabbit | Polyclonal | in-house | - | IP | - | 5 µL |
| GAD65 | Rabbit | Polyclonal | Sigma-Aldrich | ABN101 | WB | 1:1,000 | - |
|  | Mouse | Monoclonal | DSHB | GAD-6 AB_2314499 | IP | - | 3 µL |
| SNAP25 | Rabbit | Polyclonal | Synaptic Systems | 111-002 AB_887790 | WB | 1:1,000 | - |
| SNAP25 (SMI 81) | Mouse | Monoclonal | BioLegend | 836303 AB_2715864 | IP | - | 3 µL |
| GFP | Rabbit | Polyclonal | Eusera | EU2 | WB | 1:5,000 | - |
|  |  |  |  |  | IP | - | 0.75 µL |
| Mouse IgM (µ chain specific) | goat | - | Rockland Immunochemicals | 92632210 | WB (secondary) | 1:5,000 | - |
| Rabbit IgG | goat | - | Rockland Immunochemicals | 926-32211 | WB (secondary) | 1:5,000 | - |

\* Volume of antibodies used for 20 µL of Dynabeads and 200 µg – 1 mg proteins

#### 2.1.3. Experimental models

##### HD mouse lines

Heterozygous YAC128 (line 53), homozygous YAC128 (line 55), homozygous YAC18 (line 60), BACHD, humanized Hu18/18 and Hu97/18, humanized Hu21/21 and Hu128/21 mice were maintained on the FVB/N background. The description of the HD mouse lines used is displayed in **Table 3**. The YAC128 model expresses two mouse Htt alleles and the full length human mHTT transgene (Slow et al., 2003). The BACHD model expresses two mouse Htt alleles and the full length human mHTT transgene (Gray et al., 2008). The two humanized Hu97/18 and Hu128/21 models express only full length human WT and mHTT transgenes and no mouse Htt complement (Southwell et al., 2017, 2013). Experiments were performed according to protocols approved by the University of British Columbia Animal Care Committee (experimental protocol 2016/2020 #A20-0107, breeding protocol #A20-0233). Homozygous YAC128 and YAC18 mice were used for the neuronal culture, to improve culture health by avoiding an overnight genotyping step. For all cohorts, approximately equal numbers of male and female mice were used. All mice were sacrificed using CO_2_ asphyxiation in accordance with UBC SOP #I012.

**Table 3.** HD mouse lines used for palmitoylation assays.

| Mouse model | Line | Reference | Promoter | Transgene product | CAG repeat length | Endogenous product |
| --- | --- | --- | --- | --- | --- | --- |
| YAC128 | 53 | (Slow et al., 2003) | Human HTT promoter and RE (24 kb upstream, 117 kb downstream) | full length human mHTT | 128 | 2 copies of 7Q Htt |
| YAC18 | 60 |  |  | 2 copies of full length human wtHTT | 18 | 2 copies of 7QnHtt |
| YAC128 | 55 |  |  | 2 copies of full length human mHTT | 128 | 2 copies of 7Q Htt |
| BACHD | LA | (Gray et al., 2008) | Human HTT promoter and RE (20 kb upstream, 50 kb downstream) | full length human mHTT | 97 | 2 copies of 7Q Htt |
| Hu18/18 | KLAY | (Southwell et al., 2013) | Human HTT promoter and RE (20 kb upstream, 50 kb downstream) | full length human wtHTT | 18 | - |
| Hu97/18 | KLAY |  |  | full length human WT and mHTT | 97/18 | - |
| Hu21/21 | KT53 | (Southwell et al., 2017) | Human HTT promoter and RE (24 kb upstream, 117 kb downstream) | full length human wtHTT | 21 | - |
| Hu128/21 | KT53 |  |  | full length human wt and mHTT | 128/21 | - |

##### Cell line culture

Control and HD patient-derived lymphoblastoid cell lines (**Table 4**) were obtained from the NIGMS Human Genetic Cell Repository at the Coriell Institute for Medical Research. All cell lines were maintained in RPMI 1640 medium containing 10% FBS, 2 mM GlutaMAX, 10 units/mL of penicillin and 10 µg/mL of streptomycin. Cells were grown in T-25 flasks incubated in an upright position with vented caps at 37°C and 5% CO_2_. Cell density was kept between 2.10^5^ and 5.10^5^ cells/mL of media.

**Table 4.**
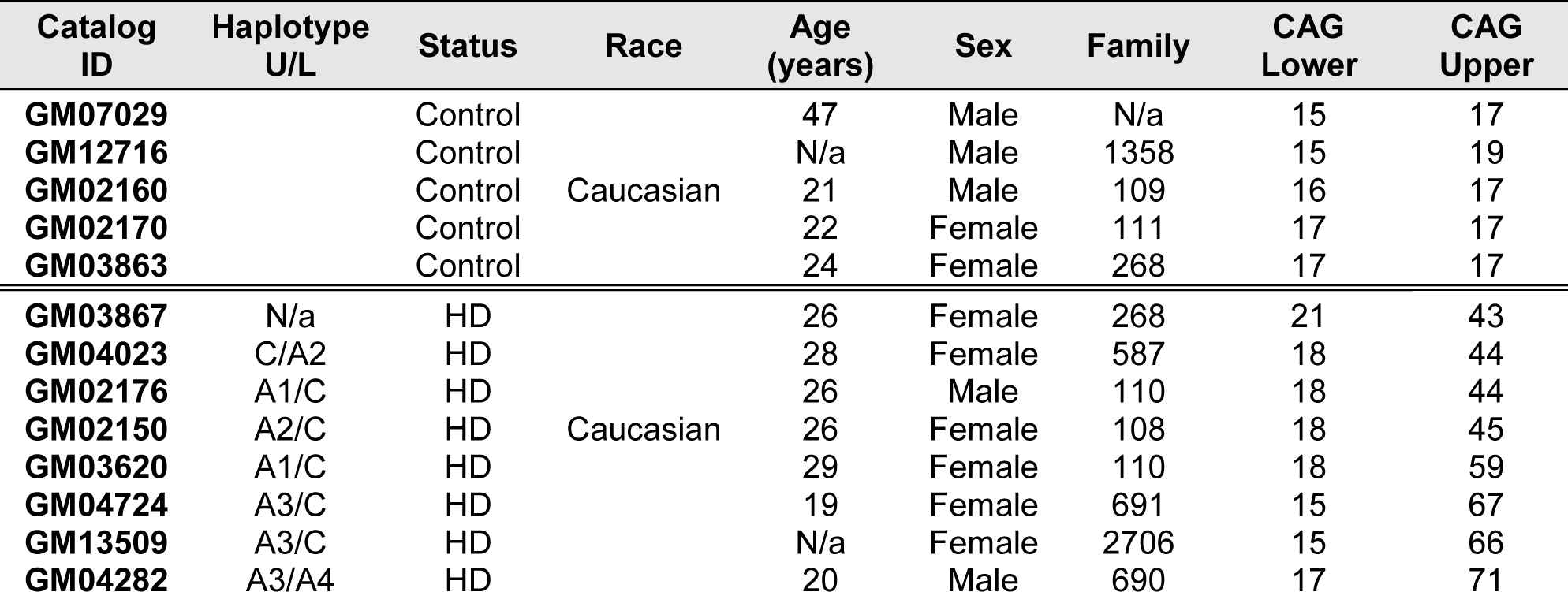
Lymphoblastoid cell lines from Coriell Biorepositories.

COS-7 cells were grown in DMEM medium supplemented with 10% FBS, 1 mM sodium pyruvate, 2 mM GlutaMAX, 10 units/mL of penicillin and 10 µg/mL of streptomycin. Cells were grown in 10-cm plates at 37°C and 5% CO_2_, and were split using a 0.25% trypsin solution (1:10 dilution) when cells were ∼90% confluent by microscopy.

Human pluripotent stem cells (hPSCs) lines listed in **Table 5** were cultured on Matrigel-coated plates in mTeSR-1 complete medium supplemented with 10 units/mL of penicillin and 10 µg/mL of streptomycin at 37°C and 5% CO_2_, and were split with ReLeSR.

**Table 5.**
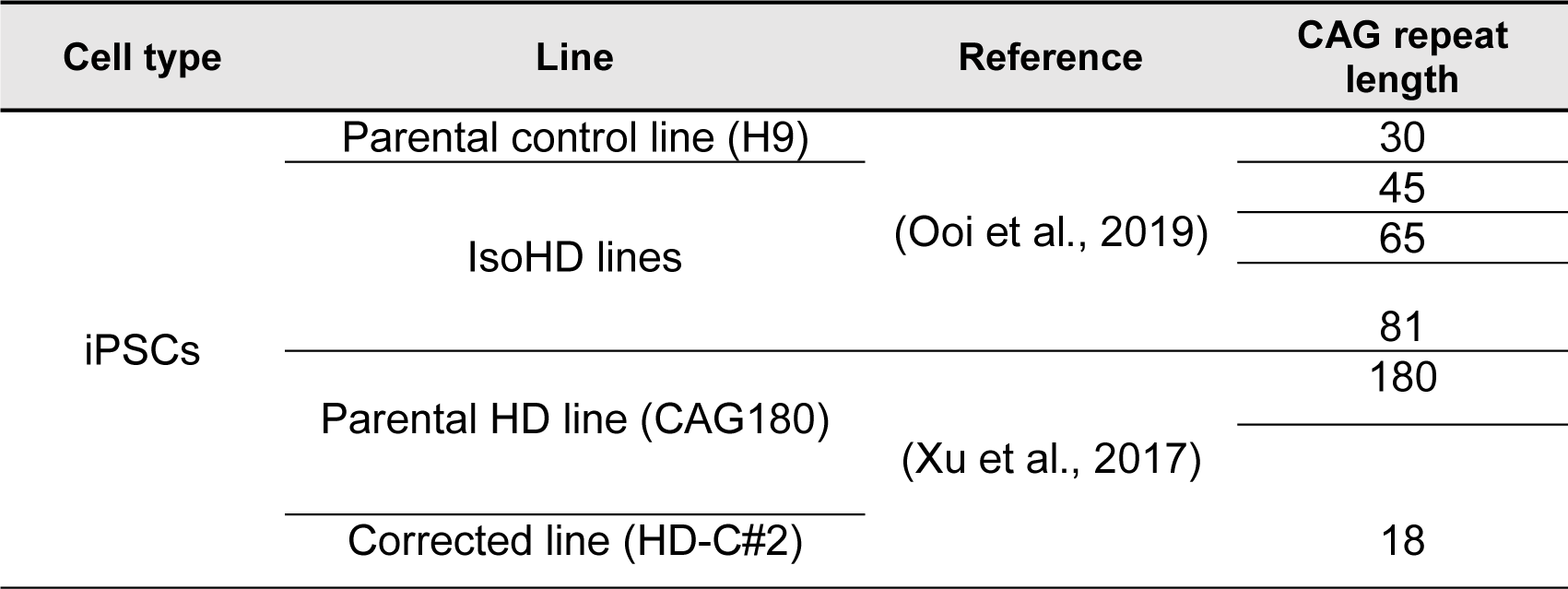
Isogenic HD patient-derived hiPSCs lines.

##### Neuronal culture

Timed pregnancies were set up by mating homozygous YAC18 (line 60) or YAC128 (line 55) mice. At E16.5, embryos were removed from pregnant females. Cortical and striatal tissues attached were dissected in ice-cold HBSS, gently dissociated with a P1000 pipette (×2-3 times) and briefly spun. Tissues were digested in 0.05% trypsin-EDTA for 8 min at 37°C until addition of 10% FBS in neurobasal (NB) medium, as previously described (Schmidt et al., 2020). Cells were centrifuged, further dissociated by mechanical trituration in NB medium (supplemented with B27, GlutaMAX, penicillin and streptomycin) and DNase (0.08 mg/mL). Neurons were plated in 6-well Nunclon Delta surface plates at a density of 1.10^6^ cells/well in 2 mL of complete NB medium. Prior to plating, 6-well plates were coated with sterile-filtered 50 μg/mL of poly-D-lysine hydrobromide for 2 h at room temperature and washed four times with sterile water. All cultures were regularly supplemented with fresh complete NB medium (20% well volume).

### 2.2. Methods

#### 2.2.1. Plasmid transfection

COS-7 cells were seeded in 6-well or 96-well plates overnight at 37°C and 5% CO_2_. The next day, cells were transfected with cDNA plasmid constructs encoding for 15Q and mutant 128Q N-terminus HTT_1-548_ (N548), HTT_1-1212_ (N1212) and full-length HTT_1-3144_ (HTT), or 17Q and 138Q HTT_1-588_-YFP (N588) using X-tremeGENE 9 DNA transfection reagent (ratio 3:1, reagent (μL): DNA (μg)). Cells were treated or harvested 18 to 24h post-transfection.

#### 2.2.2. Acyl-protein thioesterase (APT) inhibition with palmostatin B

##### Inhibition of APTs in immortalized cell lines

At 18-24h post-transfection, COS-7 cells were deprived of lipid for 1 h in DMEM (without methionine or cysteine) supplemented with 5% Charcoal stripped FBS, subsequently treated with DMSO or 15 µM Palmostatin B (PalmB) (diluted in DMSO at 1000X on InSolution) while concurrently metabolically labelled with 100 μM alkyne-palmitate for 6h. Bio-orthogonal click chemistry of alkyne-palmitate was performed as subsequently described in section §2.2.4.

##### Inhibition of APTs in primary neurons

Primary neurons from cortex and striatum of E16.5 mouse embryos (homozygous YAC18 and YAC128 mouse lines) were cultured as previously described in section §2.1.3. At DIV10, neuronal cultures were treated with DMSO or with 15 µM PalmB (InSolution) for 6h. The IP-ABE assay was performed on neuronal lysates as described in section §2.2.4.

##### Inhibition of APTs in control and HD patient-derived lymphoblastoid lines

Lymphoblasts were cultured as previously described in section §2.1.3. Cells were seeded at a density of 600,000 cells/2 mL/well in 6 well plates. The next day, cells were treated with DMSO or 10 µM PalmB for 4h. The IP-ABE assay was performed on lymphoblasts as described in section §2.2.4.

#### 2.2.3. Intracerebroventricular HTT antisense oligonucleotide infusion

A fully phosphorothioate modified gapmer ASO with 5’ locked nucleic acid (LNA) modifications in each wing, matching a sequence in intron 22 of the *HTT* gene (5’-TAATACGTAAGTGTCACAA-3’, custom synthesized by Qiagen) was delivered by intracerebroventricular (ICV) injection as previously described (Southwell et al., 2015). Five-month-old YAC128 (line 53) mice were anesthetized using 3% isoflurane and secured on a stereotaxic frame. Mice were given pre-operative subcutaneous buprenorphine (0.1 mg/kg) and 0.25% bupivacaine at the site of incision. A midline incision was made in the scalp to expose the skull. Then, a 50 µL Hamilton syringe with a 26-gauge needle was oriented to the bregma and moved 0.3 mm anterior, 1 mm lateral, and 3 mm ventral. An injection volume of 10 µL was delivered into the lateral ventricle at a rate of 5 µL/min. Mice were either injected with 50 µg ASO in sterile PBS (n=7) or PBS only (n=6). The needle was left in place for 2 min after injection, before being slowly retracted. The scalp was closed with sutures and Vetbond tissue adhesive. Animals were then placed in a recovery cage with a heating pad, hydrogel, and food on the cage floor. Mice were given post-operative buprenorphine as needed and monitored for 72h.

#### 2.2.4. Palmitoylation assays

##### Acyl-biotin exchange assay on immunoprecipitated proteins (IP-ABE)

###### Mouse brain

IP-ABE assays were performed on whole or half frozen brain samples from YAC128 (line 53) and WT littermate controls, BACHD and WT littermate controls, Hu97/18 and Hu18/18, Hu128/21 and Hu21/21, as previously described (Drisdel and Green, 2004; Huang et al., 2009). Briefly, brains were harvested and immediately snap-frozen in liquid N_2_ and stored at -80°C until being processed. Frozen brains were homogenized on ice in lysis buffer (150 mM NaCl, 50 mM Tris, 5 mM EDTA, 0.1% SDS, 1% triton X-100, pH 7.4) with 100 mM NEM (4 mL lysis buffer/brain hemisphere). Homogenates were sonicated for 15 sec at 30% power to shear DNA and the insoluble material was removed by centrifugation at 20,000 × g for 15 min at +4°C. Protein concentrations in lysates were assessed by DC protein assay. HTT, PSD95, and GAD65 were immunoprecipitated (IP) from brain lysates (300 µL) by overnight incubation with Protein G Dynabeads or PureProteome protein A/G, and appropriate antibodies for immunoprecipitation (**Table 2**). Beads were then washed and split into two and treated either with neutral pH hydroxylamine (HAM+) in lysis buffer or just lysis buffer (HAM-) for 2h for HTT and 1h for the other three proteins at room temperature. Following HAM treatment, beads were washed and treated with 2.5 µM EZ-Link BMCC-Biotin in pH 6.2 lysis buffer for 1h at 4°C. At the end of the BMCC-Biotin treatment, beads were washed and heated at 70°C for 10 min with NuPAGE LDS sample buffer and 100 mM DTT to elute proteins.

###### Control and HD-patient derived lymphoblastoid and hiPSC lines

Lymphoblastoid aggregates in suspension in the media were dissociated by trituration and centrifuged for 5 min at 500 × g at room temperature. HiPSCs were washed with DMEM/F12, lifted from plated using ReLeSR and spun at 500 × g for 5 min at +4°C. Supernatant was discarded and cell pellets were frozen until being processed, as previously described for mouse brain samples. HTT was immunoprecipitated from cell lysates by overnight incubation with Protein G Dynabeads and HTT (MAB2166) antibodies (approx. 250 µg protein and 2 µL MAB2166 for 20 µL beads).

###### Bio-orthogonal labelling assay using alkyne-palmitate

##### Immortalized cell lines

cells transiently transfected with cDNA HTT constructs were deprived of lipid during 1 h incubation in DMEM supplemented with 5% charcoal stripped FBS 18h post-transfection and subsequently metabolically labelled with 100 μM alkyne-palmitate for 6 h. Cells were lifted from plates in PBS using cell scrapers, pelleted by centrifugation (500 × g, 5 min, +4°C) and subsequently lysed in RIPA (150 mM NaCl_2_, 5 mM M EDTA, 50 mM Tris, 1% NP-40, 0.5% sodium deoxycholate, 0.1% SDS in dH_2_O, complete protease inhibitors cocktails) or SDP lysis (50 mM Tris pH 8.0, 150 mM NaCl, 1% Igepal, 40 mM NaF, 1 mM PMSF, 2 mM sodium vanadate, complete protease inhibitors cocktails) buffer for 10 min on ice, 5 min at 4°C on rotator and were centrifuged (16,000 × g, 10 min, 4°C). Bio-orthogonal click chemistry of alkyne-palmitate was performed, as previously described (Yap et al., 2010) on cell lysates immunoprecipitated with anti-HTT antibodies (BKP1 or MAB2166). HTT immunoprecipitates were adjusted to 1% SDS and incubated with 100 mM TBTA, 1 mM CuSO4, 1 mM TCEP and 100 mM azido-biotin at +37°C in darkness for 1 h.

#### 2.2.5. Western blot analysis

Brain, COS-7 and HD patient-derived hiPSC and lymphoblastoid supernatants from HTT assays were run on NuPAGE™ 3-8% Tris-Acetate gradient protein gels in Tris-Acetate SDS Running Buffer (50 mM Tricine, 50 mM Tris Base, 0.1% SDS, pH 8.24) or homemade 8% low bis acrylamide (1:200 bis-acrylamide:acrylamide) gels in SDS-Tris-glycine running buffer (25 mM Tris base, 190 mM glycine, 3.5 mM SDS) (Carroll et al., 2011) to resolve full length HTT alleles with small ΔQs. Brain lysates from the other proteins (SNAP25, PSD95 and GAD65) were run on homemade tris-glycine gels (7-12%) in SDS-Tris-glycine running buffer. Protein were transferred to 0.45 µm nitrocellulose. Membranes were blocked with 5% milk in Phosphate-Buffered Saline (PBS) or 3-5% BSA in Tris-Buffered Saline (TBS) supplemented with 0.1-0.5% Tween-20 (T). Primary antibody dilutions of HTT (D7F7 or MAB2166), SNAP25 polyclonal, PSD95 monoclonal, and GAD65 polyclonal in 3-5% BSA PBST or TBST were applied to the immunoblots at room temperature for 1h or overnight at +4°C (**Table 2**). Membranes were washed 4 × 5 min in PBST or TBST. The appropriate secondary antibodies and Alexa Fluor 680 conjugated streptavidin were applied in 3-5% BSA PBST or TBST for 1 to 2h at room temperature. Membranes were washed 4 × 5 min in PBST or TBST and imaged using the LI-COR Odyssey Infrared Imaging System (Li-Cor Biosciences). Densitometry was quantified using the LiCor Image Studio Lite software and median signal intensity following background subtraction was used for analysis. Palmitoylation was analyzed as a ratio of HAM+ palmitoylation signal to total immunoprecipitated protein signal and normalized to the control.

#### 2.2.6. Aggregation and cytotoxicity assays

##### Filter retardation assay

Protein aggregation of insoluble HTT_1-588_-YFP transiently expressed in COS-7 cells was measured using the filter-trap retardation assay, as previously described (Dale D. O. Martin et al., 2018; Wanker et al., 1999). After 72 h of transfection, COS-7 cells expressing the indicated HTT_1-588_-YFP constructs and treated with 15 µM PalmB for 48h were lysed in lysis buffer (50 mM Tris pH 8.8, 100 mM NaCl, 5 mM MgCl_2_, 0.5% Igepal CA630, 1 mM EDTA with 1X cOmplete protease inhibitor cocktail and 25 unit/mL benzonase fresh) (∼150 µL/cell pellet) for 30 min on ice without vortexing or spinning (only pipetting to homogenize). Protein concentrations in lysates were assessed by DC protein assay and adjusted to 1 µg/µL in a total volume of 50 µL. Samples (50 µg of proteins) were denatured in a 1:1 volume of denaturation buffer (2% SDS and 50 mM DTT) at 98°C for 10 min. Denatured samples were loaded in duplicate over a 0.2 µm cellulose acetate (OE66) membrane positioned in the filter retardation apparatus (Whatman, The Convertible filtration manifold system cat. Series 1055). The membrane was washed with a 0.2% SDS solution once prior to loading the samples, and twice after (100 µL/wash). The membrane was blocked in 3% BSA TBST (0.1% Tween 20) for an hour and probed by standard immunoblotting analysis with primary mouse anti-HTT (MAB2166) and rabbit anti-GFP antibodies, followed by secondary antibodies and detection using the LI-COR Odyssey immunoblotting system. The quantity of aggregates retained on the filter expressed was measured with Image Studio Lite and calculated relatively to the PBS control group (arbitrarily set at 1) and technical replicates were averaged.

##### LDH activity cytotoxicity assay

Cytotoxicity was assessed in COS-7 transiently expressing human HTT or YAC128 neurons after 6 and 24h of PalmB treatment, using the CYQUANT LDH cytotoxicity assay kit, as per the manufacturer’s guidelines. Lactate dehydrogenase (LDH) is a well-defined and reliable indicator of cytotoxicity. Extracellular LDH was here quantified by a coupled enzymatic reaction in which LDH catalyzes the conversion of lactate to pyruvate via NAD+ reduction to NADH. This can be measured by using an excitation of 560 nm and an emission of 590 nm. A cell density of 10,000 cells/well was chosen after experimental optimization. Fluorescence was read on an EnSpire 2300 Multilabel plate reader from PerkinElmer using the EnSpire manager software. Cytotoxicity (in %) was calculated using the formula:

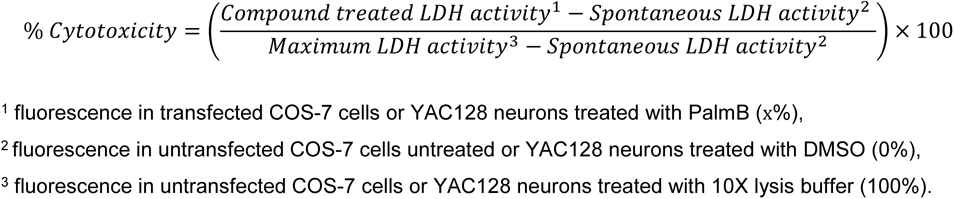

#### 2.2.7. Statistical Analysis

GraphPad Prism 9 was used for all statistical analysis and graph preparation. Figures were generated in Adobe Illustrator 2020. All data are presented as mean ± SEM. Biological replicates (n) are displayed on each graph, or indicated in the captions. Student’s t-test or 1- or 2-way ANOVA statistical tests with post-hoc analysis (Tukey, Dunnett, Sidak or Bonferroni) were used for all experiments. In some cases, t-tests were performed between specific groups after an ANOVA test when a planned comparison was established during the experimental design process. Significance for these tests is noted with the “#” symbol instead of “*”.

## 3. Results

### 3.1. Palmitoylation levels of HTT and HIP14/HIP14L substrates are decreased in the BACHD and in two humanized HD mouse models

We first wanted to assess in a low throughput, high confidence manner how the presence of mHTT impacts protein palmitoylation in multiple mouse models of HD, including BACHD, YAC128, Hu97/18 and Hu128/21 (**Table 3**). We therefore compared levels of palmitoylation of HTT as well as HIP14 and/or HIP14L substrates: 25 kDa synaptosomal-associated protein (SNAP25), 95 kDa post-synaptic density protein (PSD95), and 65 kDa glutamate decarboxylase (GAD65) in the brains of 6-month-old HD mouse lines and their respective control line (**Figure 1**).

**Figure 1.**
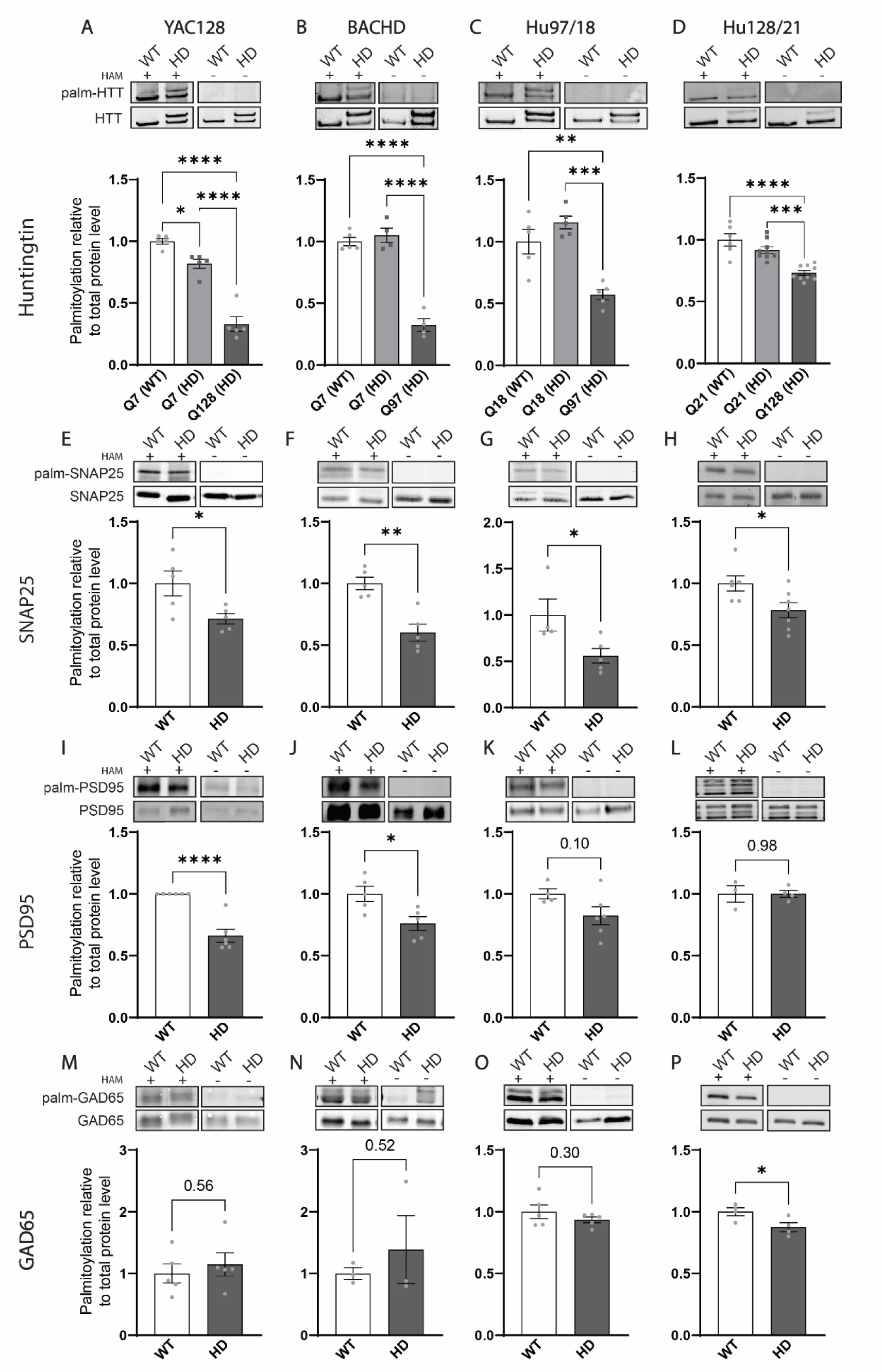
Palmitoylation levels of mutant HTT, SNAP25, PSD95 and GAD65 are reduced in the BACHD, and humanized Hu97/18 and Hu128/21 mouse models of HD. Brain lysates from 6-month-old YAC128 and littermate controls (WT), BACHD and WT, humanized Hu97/18 and Hu18/18, Hu128/21 and Hu21/21 mice were subjected to the IP-ABE assay. The palmitoylation signal (HAM+) is shown in the top panels for each mouse line and protein, and the total protein immunoprecipitated is presented in the corresponding bottom panel. The negative control (HAM-) treatment is displayed on the right of each set of blots. In the huntingtin (HTT) blots (**A-D**), the lower band corresponds to Htt or wtHTT, and the upper band to mHTT. The palmitoylation levels for HTT (**A-D**), SNAP25 (**E-H**), PSD95 (**I-L**) and GAD65 (**M-P**) were calculated as a ratio of palmitoylation over total protein signal. Mutant HTT was quantified separately from Htt or wtHTT and is represented by dark grey bars on the graphs, while Htt/wtHTT from the same HD brain is in light grey. <u>Statistical analysis</u> 1-way ANOVA: (**A**) YAC128 and (**B**) BACHD, p<0.0001; (**C**) Hu97/18, p=0.0002; (**D**) Hu128/21, p=0.0013. Stars indicate Tukey’s multiple comparisons test results. Student’s T-test: (**E-P**) results are indicated by stars or p-values on the graphs.

In the BACHD model (**Figure 1.B**), the palmitoylation level of mHTT Q97 was significantly lower (70%) than palmitoylation of mouse Htt Q7 in the WT or HD brains (1-way ANOVA (1WA): p<0.0001; Tukey’s post-test (Tk): p<0.0001 for Q97 vs. Q7 in WT or HD brains). In the humanized Hu97/18 and Hu18/18 brains (**Figure 1.C** - 1WA: p=0.0002), the palmitoylation level of mHTT Q97 was significantly lower than wtHTT Q18 in the Hu97/18 (50%; Tk: p=0.0002) and Hu18/18 (40%; Tk: p=0.0023) brains. In the Hu128/21 and Hu21/21 brains (**Figure 1.D** -1WA: p<0.0001), the palmitoylation level of mHTT Q128 was significantly lower than wtHTT Q21 in the Hu128/21 (20%; Tk: p=0.0004) and Hu21/21 (30%; Tk: <0.0001) brains. In the YAC128 model (**Figure 1.A** - 1WA: p<0.0001), palmitoylation of mHTT Q128 was significantly decreased compared to endogenous mouse Htt Q7 expressed in the YAC128 brain (60%; Tk: p<0.0001) and in the WT littermate control brain (70%; Tk: p<0.0001), consistent with the 80% decrease of mHTT Q128 compared to wtHTT Q18 in the YAC18 brain (Yanai et al., 2006).

HIP14 (ZDHHC17) and HIP14L (ZDHHC13) show substrate specificity for neuronal proteins, including SNAP25, PSD95, and GAD65 (Huang et al., 2004), which all play important roles in synaptic transmission. We therefore assessed palmitoylation levels of these proteins in all four HD mouse models. SNAP25 palmitoylation was decreased by 30% in the YAC128 brain (**Figure 1.E** - Student’s test, p=0.031), 40% in the BACHD brain (**Figure 1.F** - p=0.0017), 45% in the Hu97/18 brain (**Figure 1.G** - p=0.041) and 20% in the Hu128/21 brain (**Figure 1.H** - p=0.029) compared to their respective controls. PSD95 palmitoylation was also significantly decreased in the YAC128 brain (**Figure 1.I** - 35%; p<0.0001) and the BACHD brain (**Figure 1.J** - 25%; p=0.020) compared to WT littermate control brains. PSD95 palmitoylation showed a trend towards being reduced in the Hu97/18 brain (**Figure 1.K** - 23%; p=0.10), but was unchanged in the Hu128/21 brain (**Figure 1.L** - p=0.98) compared to controls. In contrast, palmitoylation of GAD65 was unchanged in the YAC128, BACHD and Hu97/18 mouse models (**Figure 1.M-N-O**), but was decreased by 15% in the brain of Hu128/21 (**Figure 1.P** - p=0.043) compared to the Hu21/21 mice.

Our data shows that palmitoylation levels of HTT and various HIP14/14L substrates are reproducibly reduced in all four HD mouse models. The absence of endogenous Htt in the humanized HD brain does not exacerbate protein palmitoylation dysregulation but may reveal subtle differences in palmitoylation dysregulation for GAD65 and PSD95.

### 3.2. Palmitoylation of mutant HTT is decreased early in the brain of the humanized HD mouse model, Hu128/21

Our next goal was to investigate if aging affects palmitoylation of HTT in the brain of humanized Hu128/21 HD mice. Palmitoylation of wtHTT and mHTT in the brain of Hu21/21 and Hu128/21 mice at 1, 3, 6 and 12 months of age was measured using the IP-ABE assay (**Figure 2.A-B-C-D**). At all the ages investigated, mHTT palmitoylation was significantly lower than the palmitoylation level of wtHTT expressed in the control brain (30-40%) or the HD brain (20-40%) (1WA: p<0.01 at 1 and 3 months, p<0.001 at 6 and 12 months). Notably, the decreased palmitoylation of mHTT was most pronounced at 6 and 12 months of age (**Figure 2.C and D**).

**Figure 2.**
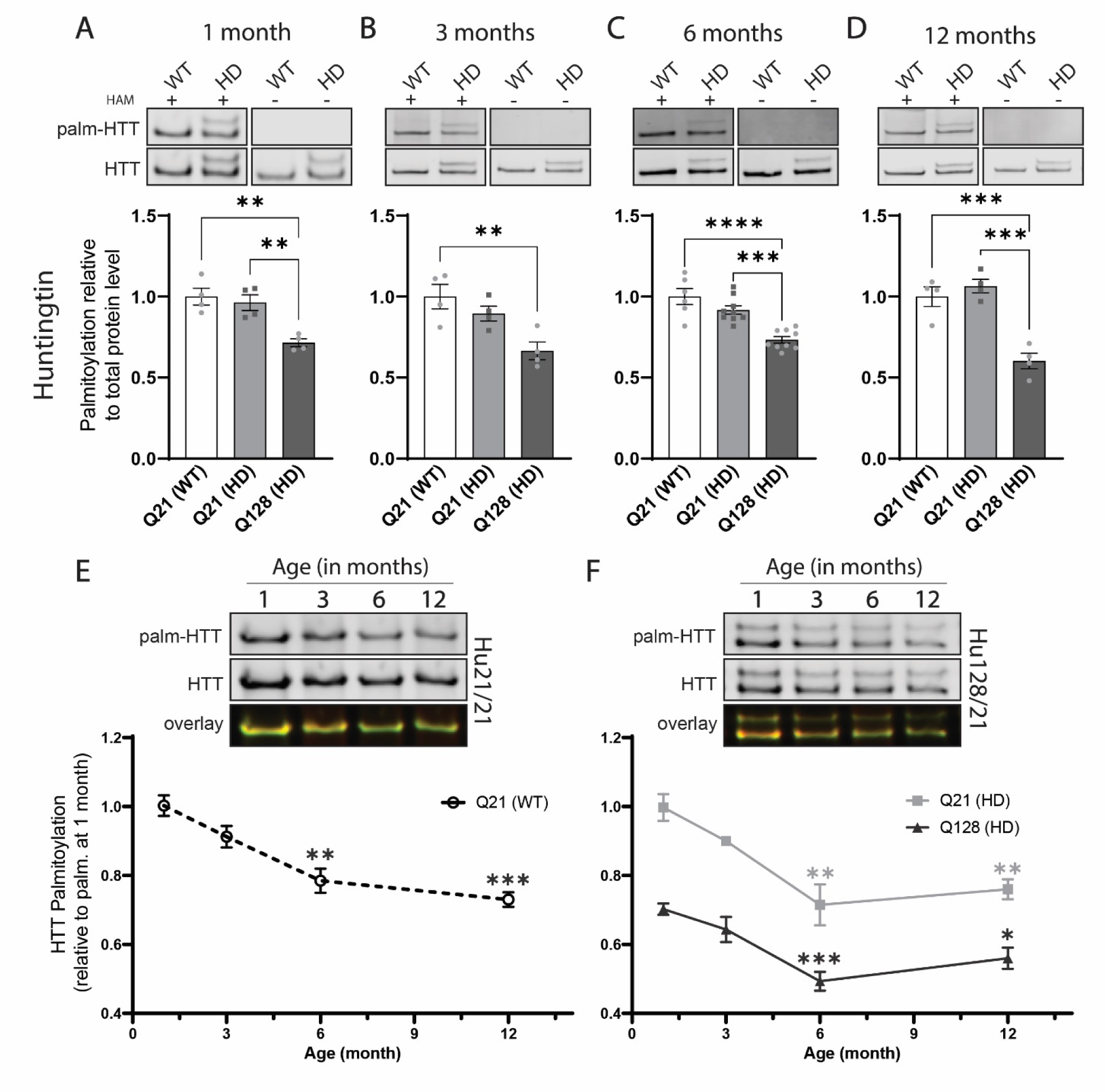
Palmitoylation of mutant HTT is decreased early in the brain of the humanized Hu128/21 mouse model of HD. HTT palmitoylation at (**A**) 1, (**B**) 3, (**C**) 6 and (**D**) 12 months measured using the IP-ABE assay in brain lysates from Hu21/21 (WT) and Hu128/21 (HD) mice. The palmitoylation signal (HAM+) is shown in the top panels for each time point, and the total protein immunoprecipitated is presented in the corresponding bottom panel. The negative control (HAM-) treatment is displayed on the right of each set of blots. Wildtype wtHTT (Q21) is the lower band and mHTT (Q128) the upper one. The quantification of 4-9 mice of each genotype is shown and was calculated as a ratio of HTT palmitoylation signal over total HTT and expressed relatively to wtHTT in the Hu21/21 brain. In the Hu128/21 brain, mHTT was quantified separately from wtHTT and is represented by dark grey bars while wtHTT from the same HD mice is represented by light bars. Effect of aging (1, 3, 6 and 12 months) on the palmitoylation levels of wt and mHTT in lysates from Hu21/21 (**E**) and Hu128/21 (**F**) brains. Total brain lysates at different ages were subjected to the IP-ABE assay and run on the same gel for each genotype (n=3-4). Overlay of both channel (palm-HTT in red and total HTT in green) are displayed in the bottom panels. Palmitoylation levels of wt and mHTT brain are expressed relatively to wtHTT palmitoylation at 1 month. <u>Statistical analysis</u>: 1-way ANOVA: (**A**) p=0.0022; (**B**) p=0.0094; (**C**) p<0.0001; (**D**) p=0.0002. Stars indicate Tukey’s multiple comparisons test results. (**E**) 1-way ANOVA: p=0.0022. Stars indicate Dunnett’s test results compared to 1 month. (**F**) 2-way ANOVA: interaction, p=0.58; age and allele effect, p<0.0001. Stars indicate Dunnett’s test results compared to wtHTT (Q21) or mHTT (Q128) at 1 month, and therefore only show the effect of age and not the effect of the HD mutation.

We then directly compared palmitoylation levels of wt and mHTT at different ages in Hu21/21 (**Figure 2.E**) or Hu128/21 brain lysates (**Figure 2.F**) and found that palmitoylation levels of wtHTT (1WA, p=0.0022) and mHTT (2-way ANOVA (2WA), p<0.0001) significantly declined with age, reaching a 30% decrease at 12 months compared to wt and mHTT palmitoylation levels at 1 month. This decline of HTT palmitoylation with age was observed for both wt and mHTT, independent of the presence of the HD mutation in the mouse brain. Taken together, our data demonstrates that palmitoylation of HTT is decreased with age and further decreased in the presence of the HD mutation as early as 1 month of age in the humanized Hu128/21 HD mouse model.

### 3.3. HTT palmitoylation levels in human lymphoblasts decrease with increasing polyQ tract length

To further characterize palmitoylation dysregulation in HD, we assessed the impact of the polyQ tract length on HTT palmitoylation in control and HD patient lymphoblasts (**Figure 3.A, Table 4**). In the control lymphoblasts, wtHTT polyQ tracts were approximately 17Qs in length. In the HD lymphoblasts, we stratified samples into two sub-groups where mHTT polyQ tract lengths were either approximately 44Q or above 59Q. The wtHTT protein in the HD lines had polyQ tracts with 17Qs.

**Figure 3.**
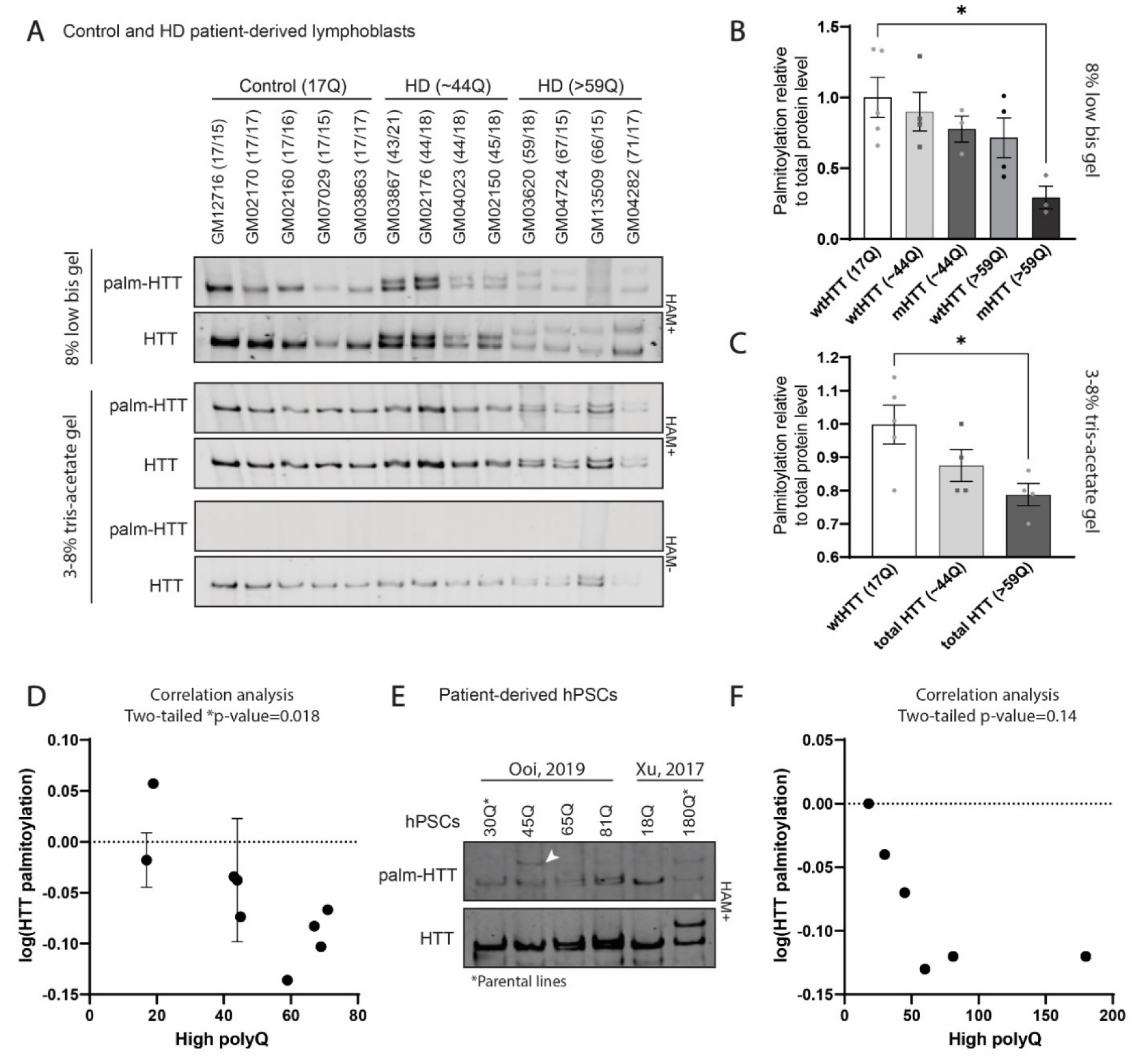
HTT palmitoylation levels decrease with increasing polyQ tract length in human lymphoblasts. (**A**) HTT (wt and mHTT) palmitoylation levels in control and HD-patient derived lymphoblasts with increasing polyQ tract length (17 to 71). Proteins were resolved using 8% low bis or 3-8% tris-acetate gels. Palmitoylation is shown in the top panels (HAM+), and the total protein immunoprecipitated with anti-HTT (D7F7) antibodies are shown in the corresponding bottom panels. The negative control (HAM-) is shown below. Each lane corresponds to a lymphoblastoid line from a control or HD patients, and represents a biological replicate (n=4-5 for each group). (**B**) Quantification of wt or mHTT palmitoylation levels in control and HD lymphoblasts in the control (17Q), HD (∼44Q) and HD (>59Q) groups after protein separation using 8% low bis gel. The quantification is shown and is calculated as a ratio of HTT palmitoylation signal over total HTT protein and expressed relatively to wtHTT palmitoylation in the control group (17Q). (**C**) Quantification of HTT (wt and mHTT together) global palmitoylation levels measured in control (17Q), HD (∼44Q) and HD (>59Q) lymphoblasts. Global palmitoylation levels of HTT was calculated as a ratio of wt and mHTT palmitoylation signal over total wt and mHTT protein and expressed relatively to wtHTT in the control (17Q) group. Mutant HTT was quantified with wtHTT, as it is challenging to resolve full length HTT alleles with small ΔQs with commonly used 3-8% tris-acetate gel. (**D**) Exponential relationship between HTT global palmitoylation levels and the polyQ tract length (high polyQ) in lymphoblasts. (**E**) HTT (wt and mutant) palmitoylation levels in HD hPSCs and isogenic controls (Ooi et al., 2019; Xu et al., 2017) with increasing polyQ tract length (18 to 180). Proteins were resolved using 3-8% tris-acetate gels. The white arrowhead indicates an unidentified band only detected with streptavidin. The hPSC parental lines used to generate HD or control hPSC lines are annotated with an asterisk. (**F**) Exponential relationship between HTT global palmitoylation level and the polyQ tract length (high polyQ) in hPSCs. <u>Statistical analysis</u>: 1-way ANOVA: (**B**) p=0.027; (**C**) p=0.037. Stars indicate Tukey’s multiple comparisons test results. (**D**) Variance explained by polyQ tract length, R^2^=0.57. Pearson r=-0.75; p=0.018. (**F**) Variance explained by polyQ tract length, R^2^=0.43. Pearson r=-0.65; p=0.15.

We quantified the palmitoylation levels of wt and mHTT separately (**Figure 3.B**) or wt and mHTT together (**Figure 3.C**) and observed that mHTT and global HTT palmitoylation levels were significantly decreased with increased polyQ tract length (1WA: (**B**) p=0.027, Tk: p=0.016 for mHTT (>59Q) vs. wtHTT (17Q) and p=0.057 for mHTT (>59Q) vs. wtHTT (∼44Q); (**C**) p=0.037, Tk: p=0.031 for total HTT(>59Q) vs. total HTT (17Q)). The exponential relationship between the global palmitoylation level of HTT (wt and mHTT together) and the polyQ tract length (high polyQ) in each lymphoblastoid line is displayed in **Figure 3.D** and supports a negative linear relationship between the two parameters (correlation, p(two-tailed) =0.018).

Palmitoylation levels were also investigated in isogenic HD human pluripotent stem cells (hPSCs) previously generated (Ooi et al., 2019; Xu et al., 2017) using the IP-ABE assay (**Figure 3.E**). The exponential relationship between the global palmitoylation level of HTT (WT and mHTT together) and the polyQ tract length (high polyQ) in each hPSC line (**Figure 3.F**) suggests a negative linear relationship between palmitoylation and polyQ length without reaching statistical significance (correlation, p(two-tailed) =0.14).

Our findings in patient-derived cell lines further validate the impact of the HD mutation on HTT palmitoylation levels and suggest that palmitoylation levels of wt and mHTT decreases with increasing polyQ length.

### 3.4 ASO-mediated mHTT lowering in the brain of the YAC128 HD mouse model does not rescue palmitoylation of HIP14 and HIP14L synaptic substrates

We next wanted to evaluate the impact of mHTT lowering in the CNS on palmitoylation of HTT and HIP14/14L substrates. Five-month-old YAC128 mice received intracerebroventricular (ICV) injection of either PBS (n=6) or an ASO targeting human HTT (HTT-ASO; n=7) (Southwell et al., 2015). Brains were collected 1 month later, and palmitoylation of HTT, SNAP25, PSD95 and GAD65 was measured using the IP-ABE assay (**Figure 4**). As expected, mHTT levels in the whole brain were lowered by approximately 90%, whereas levels of mouse Htt remained unchanged (**Figure 4.A**).

**Figure 4.**
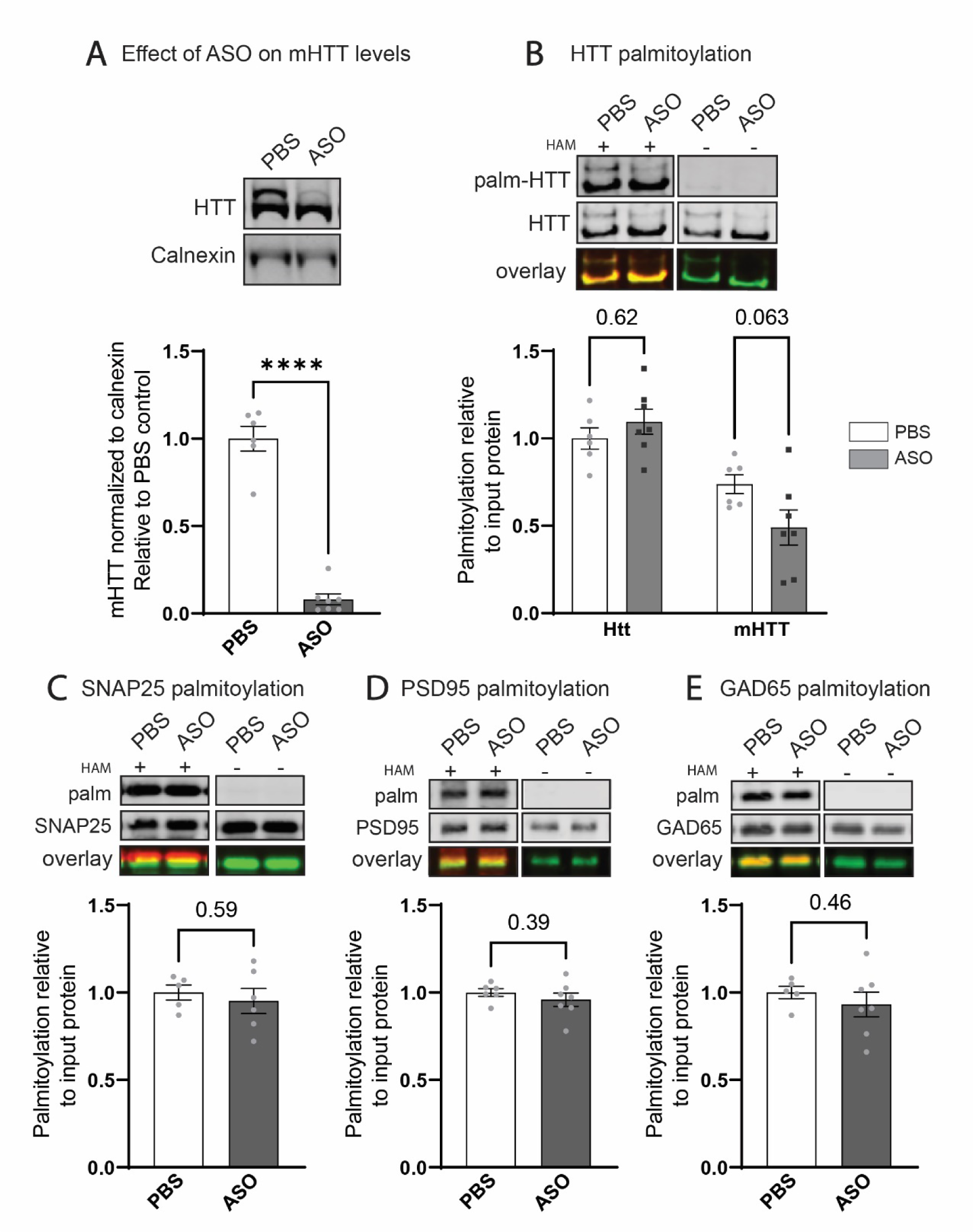
ASO-mediated mHTT lowering does not rescue aberrant palmitoylation of HIP14/HIP14L synaptic substrates in the brain of the YAC128 mouse model. Five-month-old YAC128 mice received intracerebroventricular (ICV) injection of either PBS (n=6) or an ASO targeting human HTT (HTT-ASO; n=7), and brains were collected 1 month later. (**A**) Protein from brain lysates were separated using 3-8% tris-acetate gradient gel and probed with HTT (MAB2166) and calnexin antibodies. HTT levels were quantified by densitometry and normalized to calnexin, and relative levels of mHTT are shown. Palmitoylation of HTT (**B**), SNAP25 (**C**), PSD95 (**D**) and GAD65 (**E**) was measured using the IP-ABE assay. The palmitoylation signal (HAM+) is shown in the top panels for each protein, and the total protein immunoprecipitated is presented in the panels below. The negative control (HAM-) is displayed on the right of each set of blots. Overlay of both channel (palmitoylated protein in red and total protein in green) are displayed in the bottom panels. In the HTT blots (**B**), the lower band corresponds to mouse Htt, and the upper band to mHTT. The palmitoylation levels were calculated as a ratio of palmitoylation over total protein signal. <u>Statistical analysis</u>: (**B**) 2-way ANOVA: interaction, p=0.059; allele, p<0.0001, ASO treatment, p=0.48. Stars indicate Tukey’s multiple comparisons test results between PBS and ASO groups. Student’s T-test: (**A**, **C**, **D**, **E**) results are indicated by p-values or stars on the graphs.

We measured a 30% reduction of mHTT palmitoylation compared to mouse Htt (2WA: p<0.0001, HD mutation, p<0.0001) in the PBS controls (Tk: p=0.049) and a 60% reduction in the ASO treated animals (Tk: p<0.0001), but the ASO treatment did not change Htt and mHTT palmitoylation levels (**Figure 4.B**). ASO treatment also did not significantly alter the palmitoylation level of SNAP25 (**Figure 4.C**), PSD95 (**Figure 4.D**) and GAD65 (**Figure 4.E**). Taken together, mHTT lowering in the brain of the YAC128 mouse did not rescue aberrant palmitoylation of the proteins examined.

### 3.5. Inhibition of APT increases palmitoylation levels of HTT in COS-7 cells, YAC128 cortico-striatal primary neurons and HD patient-derived lymphoblasts

Since ASO-mediated mHTT lowering *in vivo* did not rescue the reduced palmitoylation observed in YAC128 brains, we hypothesized that promoting this modification pharmacologically may represent a relevant therapeutic approach for HD. Therefore, our next objective was to assess if HTT palmitoylation levels can be upregulated in various HD models using alternative approaches. To test this, we chose to reduce protein depalmitoylation via the inhibition of the APT enzymes that catalyze thioester hydrolysis of palmitoylated cysteine residues (**Figure 5**). We chose the well characterized and membrane-permeable chiral acyl-β-lactone Palmostatin B (PalmB) (IC_50_ = 5.4 nM) that inhibits the APT1 and APT2 enzymes (Dekker et al., 2010; Vujic et al., 2016). PalmB has also been shown to partially inhibit the activity of FASN, PNPLA6, and ABHD proteins (Lin and Conibear, 2015a).

**Figure 5.**
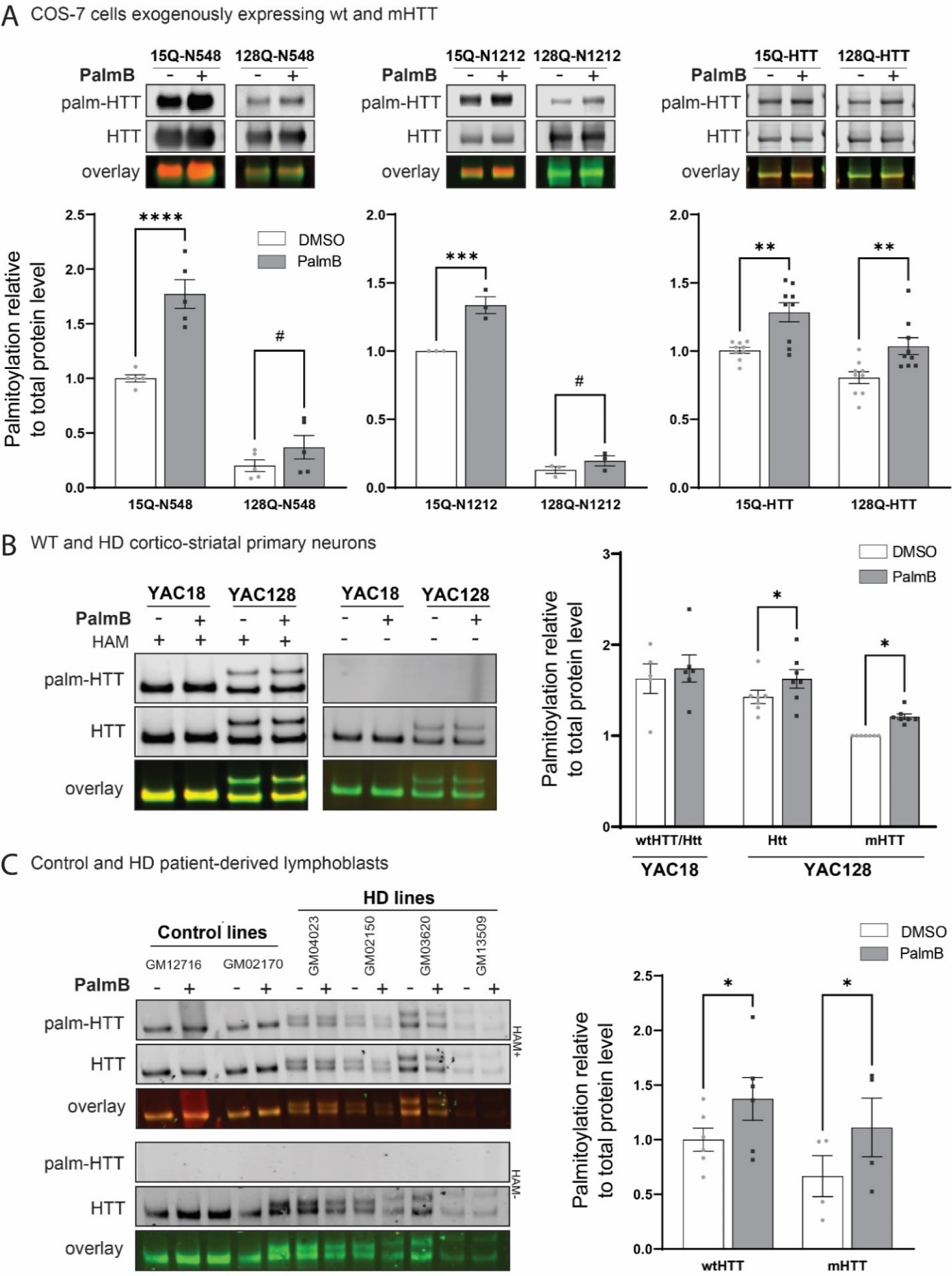
Inhibition of APTs with Palmostatin B increases palmitoylation level of HTT in COS-7 cells, YAC128 primary neurons and HD-patient derived lymphoblasts. (**A**) COS-7 cells were transfected with the N-terminal fragments HTT_1-548_ (N548), HTT_1-1212_ (N1212) and full length wt (15Q) or mutant (128Q) HTT_1-3144_ constructs. Cells were treated with DMSO or 15 μM PalmB for 6h while concurrently labelled with 100 μM alkyne-palmitate. Bio-orthogonal click chemistry of alkyne-palmitate was performed on lysates immunoprecipitated with anti-HTT (MAB2166) antibodies. Palmitoylated and total HTT were detected by immunoblots using streptavidin and anti-HTT (HD46) antibodies, respectively. Overlay of both channel (palm in red and total protein in green) are displayed in the bottom panels. Palmitoylation was calculated as a ratio of palmitoylation signal over total HTT protein and are expressed relative to wtHTT palmitoylation in the DMSO control group. (**B**) YAC18 and YAC128 cortico-striatal primary neurons were treated at DIV10 with DMSO or 15 µM PalmB for 6h. Cell lysates were subjected to the IP-ABE assay to measure palmitoylated HTT. HTT was immunoprecipitated with anti-HTT (MAB2166) antibodies and detected with D7F7 for Htt (mouse) or HTT (human) and streptavidin for palmitoylated protein. HTT palmitoylation level was calculated as a ratio of palmitoylation signal over total HTT protein and expressed relative to mHTT palmitoylation in the DMSO control group. (**C**) Control and HD-patient derived lymphoblastoid lines were treated at with DMSO or 10 µM PalmB for 4h, and lysates were subjected to the IP-ABE assay. HTT was immunoprecipitated with anti-HTT (MAB2166) antibodies and detected with D7F7 for total HTT and streptavidin for palmitoylated HTT. HTT palmitoylation level was calculated as a ratio of palmitoylation signal over total HTT protein and expressed relative to wtHTT palmitoylation level in the DMSO treated control group. <u>Statistical analysis</u>: 2-way ANOVA: (**A**) N548, interaction, p=0.0042; HD mutation, p<0.0001; PalmB, <0.0001. N1212, interaction, p=0.0075; HD mutation, p<0.0001; PalmB, p=0.0007. Full length HTT, interaction, p=0.65; HD mutation, p=0.0002; PalmB, p<0.0001. (**B**) Interaction, p=0.86; HD mutation, p<0.0001; PalmB, p=0.0320. (**C**) Interaction, p=0.90; HD mutation, p=0.43; PalmB, p=0.0071. Stars indicate Bonferroni’s multiple comparisons test results and hashtag t-test results.

The impact of PalmB treatment on mHTT palmitoylation levels in COS-7 cells transiently expressing N-terminal fragments HTT_1-548_ (N548), HTT_1-1212_ (N1212) or full length HTT_1-3144_ 18Q (wt) or 128Q was assessed by metabolic labelling with palmitic acid alkyne followed by bioorthogonal labelling (click chemistry) (**Figure 5.A**). The PalmB concentration of 15 µM was selected after testing a wide range of concentrations (0, 10, 15, 20 and 50 µM) in our experimental models. Palmitoylation levels of N548, N1212 and full length HTT were significantly decreased in the presence of the HD mutation, with a stronger effect for shorter fragments (2WA: HD mutation, p<0.0001 for N548 and N1212, and p=0.0002 for full length HTT). Treatment of COS-7 cells with 15 µM PalmB for 6h significantly increased the palmitoylation levels of 15Q and 128Q N548, N1212 and full length HTT compared to the respective DMSO controls (2WA: PalmB effect: p<0.0001 for N548 and full length HTT, and p=0.0007 for N1212). Notably, palmitoylation levels of full length mHTT were restored to those of wtHTT following treatment with PalmB.

We next used primary cortico-striatal neuronal cultures from YAC18 and YAC128 mice to evaluate the impact of PalmB treatment on HTT palmitoylation levels (**Figure 5.B**). As expected, the palmitoylation level of mHTT was significantly decreased compared to mouse Htt and wtHTT (2WA: HD mutation, p<0.0001). Treatment with PalmB significantly increased human HTT palmitoylation (2WA: PalmB effect, p=0.032). In the YAC128 neuronal cultures, mouse Htt and mHTT palmitoylation were significantly increased by the PalmB treatment compared to the DMSO control by approximately 20% (Bonferroni’s test (Bf): p=0.041 and p=0.028). In the YAC18 neuronal culture, the effect of PalmB on the palmitoylation levels of mouse Htt and human wtHTT (not distinguishable with our protocol) was weaker (Bf: p=0.54).

Finally, we explored the impact of PalmB treatment on HTT palmitoylation in control and HD-patient derived lymphoblasts (**Figure 5.C**). Notably, treatment with PalmB significantly increased HTT palmitoylation in this model (2WA: PalmB effect, p=0.0071), rescuing mHTT palmitoylation levels to wtHTT palmitoylation level.

Inhibiting the activity of the APT enzymes with PalmB treatment significantly increased full length HTT palmitoylation levels in immortalized cell lines, primary YAC128 neuronal cultures and HD patient-derived cells. Therefore, the impact of the HD mutation on mHTT palmitoylation is reversible and can be modulated using a small molecule APT inhibitor.

### 3.6. Inhibition of APTs with PalmB treatment reduces mHTT-induced cytotoxicity in COS-7 cells and YAC128 primary neurons

Our last objective was to investigate if increasing mHTT palmitoylation levels through APT inhibition would impact mHTT inclusion formation and/or mHTT-induced cell toxicity in an *in vitro* HD model. We measured the impact of PalmB treatment in COS-7 cells transfected with mutant 138Q HTT_1-588_-YFP (138Q-N588-YFP) on the quantity of insoluble mHTT inclusions using a filter retardation assay (**Figure 6**). We confirmed that PalmB treatment increased 138Q-N588-YFP palmitoylation level (by 30%) compared to the DMSO treated control (p=0.029) (**Figure 6.A**), and that the amount of insoluble HTT inclusions were significantly higher following expression of 138Q-N588-YFP compared to 17Q-N588-YFP (p<0.0001) (**Figure 6.B**). Treatment with PalmB for 48h significantly decreased the formation of insoluble inclusions by 20-22% (p=0.038 and 0.0065 with HTT or GFP antibodies) (**Figure 6.C**).

**Figure 6.**
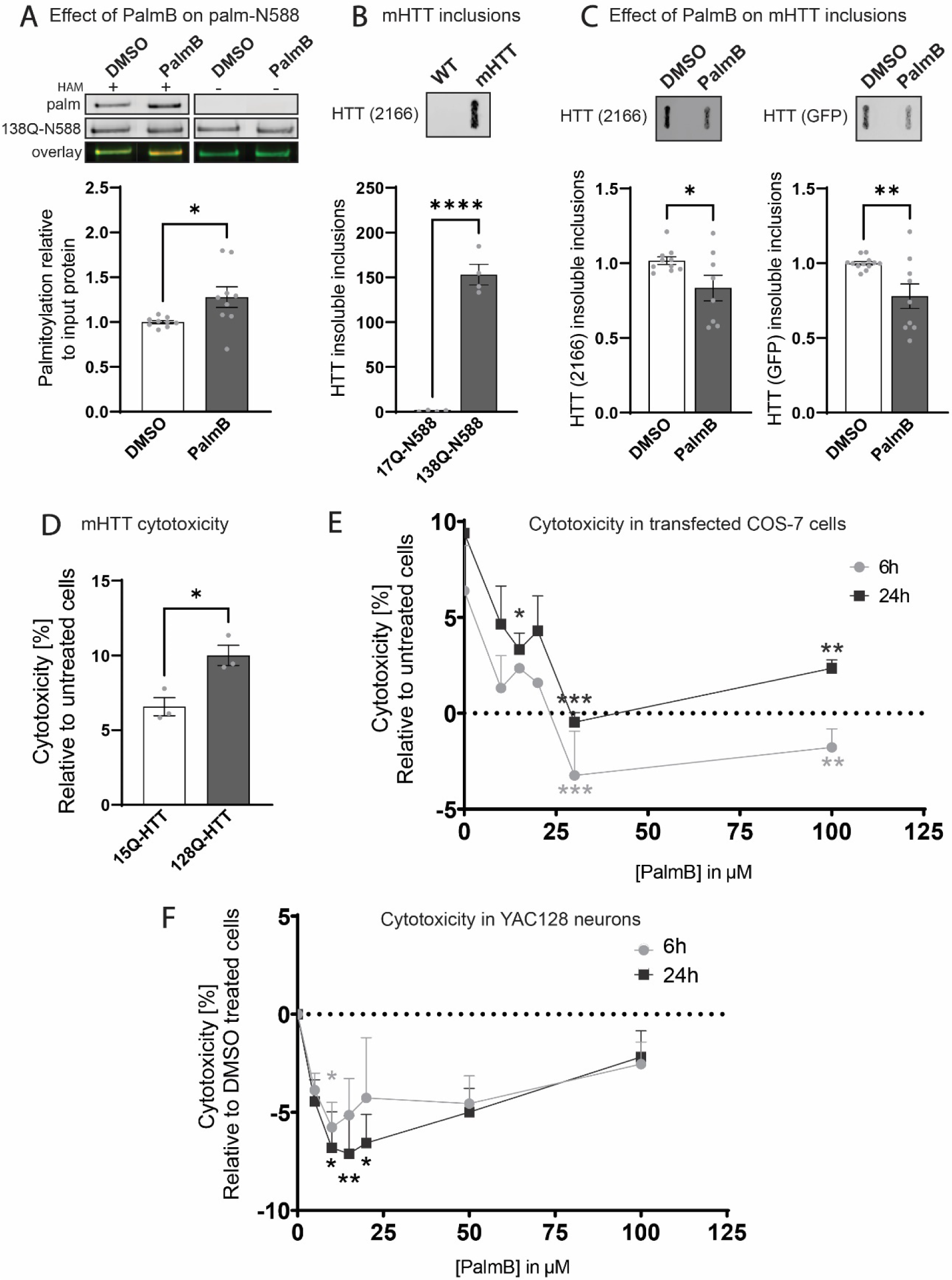
Inhibition of APTs with Palmostatin B reduces mHTT insoluble inclusions in COS-7 cells and mHTT-induced cytotoxicity in COS-7 cells and YAC128 cortico-striatal primary neurons. COS-7 cells were transfected at t=0h with 138Q-HTT_1-588_-YFP (138Q-N588-YFP) construct, treated with 15 µM PalmB at t=24h and 48h post-transfection and harvested at t=72h. (**A**) Cell lysates were subjected to the IP-ABE assay to measure palmitoylated HTT. HTT was immunoprecipitated with anti-GFP antibodies and detected with MAB2166 for HTT and streptavidin for palmitoylated protein. HTT palmitoylation level was calculated as a ratio of palmitoylation signal over total HTT protein and expressed relative to palmitoylation in the DMSO control group. (**B-C**) Insoluble mHTT inclusions were monitored by filter retardation assay and immunodetection with anti-HTT (MAB2166) or GFP antibodies. The amount of aggregates retained on the filter is expressed relative to (**B**) 17Q-N588-YFP or (**C**) to the DMSO treated group. (**D**) Expression of full length 128Q-HTT in COS-7 cells increased cytotoxicity compared to 15Q-HTT expression. COS-7 cells seeded in 96-well pates were transfected with WT (15Q) or mutant (128Q) full length HTT for 24h. LDH activity in conditioned media was measured using the CyQUANT cell proliferation assay. Cytotoxicity (%) is expressed relative to untreated and non-transfected cells. (**E**) Cytotoxicity in COS-7 cells transfected for 24h with full length 128Q-HTT and treated with DMSO or PalmB (InSolution at 10, 15, 20, 30 and 100 µM) for 6 or 24h was assessed using the LDH activity assay (n=3). Cytotoxicity (%) is expressed relative to untreated and non-transfected cells. Negative cytotoxicity values correspond to a protective effect. (**F**) Cytotoxicity in YAC128 cortico-striatal primary neurons at DIV15 treated with DMSO or PalmB (InSolution at 5, 10, 15, 20, 50 and 100 µM) for 6 or 24h, using the LDH activity assay (n=4). Cytotoxicity (%) is expressed relatively to YAC128 neurons treated with DMSO. Negative cytotoxicity corresponds to a protective effect. Negative cytotoxicity values correspond to a protective effect. <u>Statistical analysis</u>: Student’s T-test (**A**) p=0.029, (**B**) p<0.0001, (**C**) MAB2166, p=0.038 and GFP, p=0.0065. (**D**) p=0.019. 2-way ANOVA: (**E**) Interaction, p=0.90, PalmB effect, p<0.0001, Treatment duration, p=0.0033; (**F**) Interaction, p=0.97; PalmB, p=0.0014; Treatment duration, p=0.29. Stars indicate Dunnett’s test results compared to respective DMSO control.

The impact of APT inhibition on cytotoxicity was measured using the LDH activity assay in COS-7 cells transiently expressing 15Q or 128Q-HTT_1-3144_. First, we confirmed that the expression of full length 128Q-HTT increased cytotoxicity (relative to untransfected cells) compared to 15Q-HTT (p=0.019) (**Figure 6.D**). Then, we measured the impact of PalmB treatment for 6 or 24h on cytotoxicity in COS-7 expressing 128Q-HTT (**Figure 6.E**). Negative cytotoxicity values correspond to a protective effect of the treatment. Treatment with PalmB for 6 and 24h significantly reduced cytotoxicity in COS-7 cells expressing 128Q-HTT relative to DMSO controls (2WA: PalmB effect, p<0.0001).

The effect of PalmB on cytotoxicity compared to the DMSO control was the strongest at a concentration of 30 µM, but was also significant at lower concentrations (Dunnett’s test: p<0.05).

The impact of PalmB treatment on cytotoxicity was also assessed in YAC128 cortico-striatal primary neurons (**Figure 6.F**). Cytotoxicity was significantly reduced by the PalmB treatment compared to neurons treated with DMSO (2WA: PalmB effect, p=0.0014). The effect of PalmB against cytotoxicity compared to the DMSO treated control was the strongest with 10 µM and 15 µM after 6 and 24h or treatment, respectively (Dunnett’s test: p=0.048 and 0.0098).

Altogether, the data shows that inhibiting of APTs with PalmB reduces mHTT inclusion levels and mHTT-induced cytotoxicity in COS-7 cells and YAC128 cortico-striatal primary neurons.

## 4. Discussion

Targeting the cause of HD, by lowering levels of the toxic mHTT protein, is one of the therapeutic strategies for the treatment of this disease (Caron et al., 2018). This can be achieved by suppressing transcription of *HTT*, degrading the transcript, reducing synthesis of the toxic protein or increasing its degradation in cells. Promising strategies to promote mHTT degradation use mHTT-LC3 linker compounds (Li et al., 2019) or proteolysis-targeting chimera (PROTAC) based approaches (Harding and Tong, 2018). Another approach has been to target specific PTMs of HTT that are dysregulated in presence of the HD mutation in order to promote neuronal survival (Aharony et al., 2015; Bowie et al., 2018; Di Pardo et al., 2012; Ehrnhoefer et al., 2011; Gu et al., 2009; Kratter et al., 2016).

The challenge with this strategy lies in the identification of the enzymes that modulate PTMs with a high specificity for HTT. Most kinases, phosphatases or caspases have a large variety of substrates. Developing therapeutically relevant pharmacological modulators of PTMs requires that they be developed within the context of well-characterized PTM-related signaling systems. However, there are still many knowledge gaps regarding the role of PTM dysregulation on mHTT toxicity in HD that need to be addressed to develop effective therapies.

A potential PTM candidate is palmitoylation of HTT at cysteine 214, catalyzed by HIP14 (ZDHHC17) and HIP14L (ZDHHC13) (Yanai et al., 2006). Here, we showed that palmitoylation levels of HTT and various HIP14/14L substrates were reproducibly reduced in a variety of HD models, including humanized HD mouse models, HD patient-derived lymphoblast and isogenic hiPSCs (**Figures 1** and **3**). Palmitoylation of HTT was decreased as early as 1 month of age in the HD humanized mouse model and palmitoylation levels were further decreased with aging (**Figure 2**). Lowering mHTT levels in the YAC128 mouse brain was not sufficient to rescue palmitoylation levels of synaptic HIP14/14L substrates (**Figure 4**). Treatment of COS-7 cells, YAC128 cortico-striatal neurons and in HD patient derived lymphoblast with Palm B, a small molecule that targets mainly APTs 1 and 2, normalized palmitoylation of full length mHTT (**Figure 5**). Finally, modulating mHTT palmitoylation was beneficial against mHTT-induced cytotoxicity (**Figure 6**).

Our first aim was to investigate how the HD mutation impacts HTT palmitoylation in multiple mouse and human HD models to address model biases and technical limitations previously encountered and to obtain a more accurate representation of palmitoylation dysregulation that may occur in HD patients. The palmitoylation of mHTT was previously shown to be reduced by 50% in COS-7 cells expressing 15Q and 128Q N548 and metabolically labelled with [^3^H]-palmitate, and by 80% in the YAC128 HD mouse brain compared to wtHTT in the YAC18 brain (a different mouse line than YAC128 expressing endogenous mouse 7Q-Htt and human 18Q-HTT) (Yanai et al., 2006) followed by the earliest version of the ABE assay (Drisdel and Green, 2004), which has since been optimized to yield higher sensitivity (Brigidi and Bamji, 2013).

A potential confounding factor of using the YAC128 and BACHD mouse models is the presence of two endogenous mouse Htt alleles in addition to the mHTT transgene, which could compensate for the effect of mHTT on HIP14 and HIP14L activity. To take this into consideration, the two humanized HD mouse models, Hu97/18 and Hu128/21, which harbor a mutant and a wtHTT human transgene with no mouse Htt complement, were also used to investigate palmitoylation in a model that more closely models the human condition. In all four mouse models of HD, mHTT palmitoylation was reproducibly decreased compared to littermate control mice and compared to the WT protein within the same mouse (**Figure 1**).

To assess HTT palmitoylation in a more physiologically relevant system, we used control and HD patient-derived lymphoblasts. We observed that palmitoylation of HTT is decreased with increasing number of polyQ repeats (**Figure 3.A-D**). This is, to our knowledge, the first demonstration of aberrant palmitoylation of mHTT in a human-derived HD cell model. This supports the finding that decreased palmitoylation of mHTT in the brains of a variety of HD mouse models is not an artifact due to lower affinity of mouse HIP14/14L (or potentially other PATs) for the human huntingtin protein, but a robust biochemical phenotype. Previous studies have looked at palmitoylation dysregulation in HD models with extreme polyQ lengths (>90) rarely seen in the clinic (Huang et al., 2011; Yanai et al., 2006). Our results now suggest that aberrant palmitoylation of HTT occurs within a range of clinically-relevant polyQ-repeat lengths found in HD patients. We also provide the first evidence that mHTT palmitoylation levels are inversely correlated with polyQ repeat length. The data generated in two sets of isogenic control and HD (IsoHD) hPSC lines (**Figure 3.E-F**) suggests that the differences of palmitoylation level are not due to additional factors such as genetic variants beyond the *HTT* gene.

Our next objective was to investigate the earliest timepoint at which palmitoylation is altered in the humanized HD mouse model, Hu128/21 (**Figure 2**). We showed that palmitoylation of mHTT was decreased very early in the whole brain (at 1 month), before onset of behavioural and neuropathological HD-like phenotypes (Southwell et al., 2017), making it one of the earliest phenotypes observed in this model. This downregulation of mHTT palmitoylation is maintained at 3, 6 and 12 months in the brain of Hu128/21. Additional studies are required to determine if mHTT is hypo-palmitoylated during embryonic development.

We report here for the first time that aging led to a decrease of both wtHTT and mHTT palmitoylation. This suggests that the presence of the HD mutation has the same impact as aging on HTT palmitoylation levels. The combined impact of aging and the HD mutation may be additive in reducing palmitoylation levels. The relationship between palmitoylation levels of proteins in aging have not been extensively investigated (Zamzow et al., 2019).

Therapeutics that lower toxic mHTT have shown promise in preclinical studies in HD rodents (Kordasiewicz et al., 2012; Southwell et al., 2018) and are currently being evaluated in clinical trials for HD (NCT04120493, NCT04000594 and NCT03225833). We have previously shown that reducing levels of Htt in *Htt*^+/−^ mice and FVB neurons treated with pan-HTT ASO (∼95% HTT knockdown) leads to reduced palmitoylation of HIP14 and HIP14 substrates such as SNAP25 and GluR1 (Huang et al., 2011). However, the impact of lowering mHTT on palmitoylation of HIP14 and HIP14L substrates has not been investigated. Our data from YAC128 animals treated with an ASO against mHTT at 5 months of age and assessed at 6 months of age (**Figure 4**; ∼90% reduction of mHTT levels in the whole brain) suggests that mHTT lowering for 1 month is not sufficient to rescue palmitoylation alterations in HD. This suggest that specific palmitoylation-targeted therapies may be necessary. Additional studies to investigate the impact of ASO treatment at a younger age, or for a longer duration are needed.

Aberrant palmitoylation causes or contributes to a large variety of disorders such as Alzheimer’s disease, schizophrenia, mental retardation and Batten Disease (Bhattacharyya et al., 2013; Cho and Park, 2016; Hofmann and Lu, 2014; Ko and Dixon, 2018). The concept of developing therapeutically relevant pharmacological modulators of palmitoylation to promote or reduce palmitoylation of specific proteins has been an emerging field over the past years (Chavda et al., 2014; Fraser et al., 2019). In this study, we observed robust reduction of HTT palmitoylation in a variety of HD models. HTT is dynamically palmitoylated by HIP14 and 14L and de-palmitoylated by APTs 1 and 2 (also LYPA1 and 2) (Lin and Conibear, 2015b; Yanai et al., 2006). We have previously shown that increasing mHTT palmitoylation levels *in vitro* by overexpression of HIP14 may be protective (Yanai et al., 2006), but this approach is not easily translatable *in vivo*. Inhibition of APTs with the membrane-permeable chiral acyl-β-lactone PalmB (Davda and Martin, 2014; Dekker et al., 2010; Vujic et al., 2016) was shown to increase HTT_1-548_ palmitoylation levels in COS-7 cells (Lin and Conibear, 2015b). Here, we demonstrate that PalmB can be used to promote palmitoylation of full length mHTT in multiple models (**Figure 5**).

This study serves as a proof-of-concept that mHTT palmitoylation can be increased, and even normalized to wtHTT levels, using a small molecule. Furthermore, we show that increasing palmitoylation of mHTT exerts beneficial effects on aggregation and cytotoxicity (**Figure 6**). Protective effects of promoting palmitoylation of HTT and other substrates may be mediated by increased mHTT clearance, or reduced proteolytic cleavage of mHTT by caspase 6 (CASP6) at D586. We have indeed recently shown that CASP6 is palmitoylated by HIP14, resulting in CASP6 inactivation (Lontay et al., 2020; Skotte et al., 2017) which would be expected to reduce cleavage and potentially provide beneficial effects.

Further studies evaluating the impact of promoting HTT palmitoylation on well characterized HD phenotypes will help elucidate the potential mechanism of action leading to the observed protective effects. PalmB inhibits APT 1 and 2 that depalmitoylate a large variety of protein substrates, and also targets FASN, PNPLA6, and ABHD proteins (Lin and Conibear, 2015a). Testing additional APT inhibitors (Vujic et al., 2016) or developing new candidate small molecules that specifically disrupt the interaction between APTs and HTT or HIP14 will be important for advancing this therapeutic strategy for HD.

## Acknowledgments & Funding

The authors would like to thank Mark Wang, Xiaofan Qu and Qingwen Xia for their technical support, Dr. Amirah Ali, Dr. Philip Ly, Hailey Findlay-Black, Jennifer Collins, Dr. Niels Skotte, Dr. Amber Southwell and Stephanie Bortnick for their support on the project. FLL was funded by a Canadian Institutes of Health Research (CIHR) Postdoctoral Fellowship. NSC was funded by the Ripples of Hope Pfizer Trainee Award, a Canadian Institutes for Health Research Postdoctoral Fellowship, and the Huntington Disease Society of America Berman/Topper Family HD Career Development Fellowship. MES was supported by a Vanier Canada Graduate Scholarship and a University of British Columbia 4-Year Graduate Fellowship. MAP is Michael Smith Foundation for Health Research Scholar. Project operational support for MRH provided by a CIHR Foundation grant (FDN-154278) and by the CHDI Foundation.

## Abbreviations

ABE: acyl-biotin exchange assay
APT: acyl-protein thioesterase
ASO: antisense oligonucleotide
HD: Huntington disease
hiPSC: human induced pluripotent stem cell
HIP14: huntingtin-interacting protein 14 (ZDHHC17)
HIP14L: huntingtin-interacting protein 14-like (ZDHHC13)
HTT: huntingtin (human)
Htt: huntingtin (mouse)
ICV: intracerebroventricular
LNA: locked nucleic acid
mHTT: mutant huntingtin
NEM: N-ethylmaleimide
PalmB: palmostatin B
PAT: palmitoyl acyltransferase
PTM: post-translational modification

